# Pleiotropic functions of glutathione in the adaptive response to long-term nitrogen starvation in *Escherichia coli*

**DOI:** 10.1101/2025.10.31.685819

**Authors:** Harriet Ellis, Volker Behrends, Gerald Larrouy-Maumus, Josh McQuail, Sivaramesh Wigneshweraraj

**Author notes:** To whom correspondence should be addressed:., Correspondence may also be addressed to Josh McQuail.

## Abstract

Nitrogen (N) is essential for bacterial growth, and adaptation to N starvation involves extensive reprogramming of gene expression. A hallmark subcellular feature in long-term N starved *Escherichia coli* cells is the presence of biomolecular condensates of the major bacterial RNA regulator Hfq. The Hfq condensates, which accumulate gradually during N starvation, contribute to adaptation by modulating RNA metabolism and central metabolic pathways. Metabolites play central roles in stress responses, often acting as modulators of protein function to support survival and recovery. Glutathione (GSH), a universal stress protectant, has broad roles in bacterial stress adaptation, yet its function during N starvation remains unexplored. Using a GSH-deficient mutant (Δ*gshAB*), we show that GSH is required for optimal survival and recovery from prolonged N starvation. We reveal that GSH regulates the temporal dynamics of Hfq condensation and dissipation during starvation and recovery, respectively, via an as-yet unknown mechanism. Notably, these two functions of GSH appear mutually exclusive, highlighting its pleiotropic role in the adaptive response to N starvation that potentially extends its canonical function as a stress protectant.

**IMPORTANCE:** Nitrogen is a vital nutrient that bacteria need to grow. When nitrogen becomes scarce, bacteria must quickly adjust how their genes are used to survive. In *Escherichia coli*, one of the key changes during long-term nitrogen starvation is the formation of tiny structures inside the cell called Hfq condensates, which help manage genetic information flow and metabolism. Small molecules called metabolites play important roles in helping bacteria cope with stress, and one such molecule, glutathione (GSH), is known to protect cells under various stress conditions. However, its role during nitrogen starvation is not known. In this study, we used a mutant strain of *E. coli* that cannot produce GSH and found that these bacteria struggle to survive and recover from nitrogen starvation. We also discovered that GSH helps control when and how Hfq condensates form and disappear. Although, these two functions of GSH seem to be unrelated, our study highlights the versatile role of GSH in helping bacteria adapt to nitrogen stress.

## INTRODUCTION

Nitrogen (N) is used for the biosynthesis of the building blocks of all proteins (amino acids), nucleic acids (nucleotides) and metabolites and cofactors in bacteria. As such, N is an essential component for bacterial growth. Notably, bacteria in the mammalian gut exist in a N-starved state as they only have access to an average of just one N atom for every ten carbon atoms (1). This is because mammals have evolved to keep homeostasis of the gut bacterial community by starving them of N. Similarly, many freshwater, marine and terrestrial ecosystems, where bacteria prevail, are limited for N (2,3). Further still, many bacterial pathogens are thought to experience N limitation in host environments such as in the urinary tract (e.g., uropathogenic *Escherichia coli*) or *Salmonella* Typhimurium (in macrophages) (4,5). The fact *E. coli* respond to N deficiency by assuming the ‘persister phenotype’ capable of evading killing by antibiotics (6) suggests a role for the adaptive response to N starvation in bacterial antibiotic recalcitrance. When bacterial systems are used for bioproduction, bacterial growth is often decoupled from bioproduction to maximise yield or metabolic pathways reprogrammed to direct production of specific biomolecules. This is often achieved by modulating N availability (7). Clearly, elucidating the mechanisms by which enteric bacteria adapt to changes in N availability is fundamental to our understanding of bacterial stress-adaptation, pathogenesis, and the rational design of bacteria for bioproduction.

Studies from many groups, including ours, have used *E. coli* as a model system to provide a detailed picture of the gene expression changes that underpin the adaptive response to N starvation. Briefly, in *E. coli* and related bacteria, N is required for the synthesis of glutamate (for protein synthesis) and glutamine (for synthesis of nucleic acids). The enzyme glutamate dehydrogenase catalyses the reductive amination of α-ketoglutarate (α-KG; a key intermediate of the Krebs cycle) to glutamate (Fig. 1A(i)). Subsequently, glutamine synthetase catalyses the amidation of glutamate to glutamine (Fig. 1A(i)). Both, glutamate dehydrogenase and glutamine synthetase use ammonium as the N source for the synthesis of glutamate and glutamine. The intracellular concentration of glutamine is the main signal for N availability in *E. coli*, and its levels are detected by the uridylyltransferase/uridylyl-removing enzyme, GlnD (for reviews see (8-10)). As shown in Fig. 1A(ii), under N replete conditions, when glutamine concentrations are high, GlnD deuridylylates GlnB. The deuridylylated form of GlnB binds to NtrB to activate its phosphatase activity and consequently dephosphorylates NtrC, thereby inactivating it. The deuridylylated form of GlnK interacts with the ammonium transporter, AmtB, to inhibit ammonium uptake. Conversely, under N starvation, when the intracellular concentration of glutamine is low, GlnB and GlnK become uridylylated, which prevents the inhibition of AmtB by GlnK, thus enabling the uptake of ammonium, and phosphorylation of NtrC (by NtrB) leading to expression of the NtrC regulon. The NtrC regulon is extensive and results in the transcriptional reprogramming of ∼40% of all *E. coli* genes (11). Emerging results have revealed that, in addition to transcriptional reprogramming, post-transcriptional regulation of RNA, mediated by the major bacterial RNA chaperone Hfq, plays a major role in the adaptive response to N starvation in *E. coli* (12-14). We discovered that, during N starvation, ∼50% of Hfq molecules in *E. coli* progressively assemble into a foci-like structure near the cell poles (15,16). We term these structures ‘condensates’ because they form through a liquid-liquid phase separation-like process and rapidly disperse upon N replenishment. Both, the formation and dispersion of Hfq condensates occur independently of gene expression, suggesting that the function of Hfq condensates extends beyond that of simply Hfq’s canonical function in facilitating the interaction between non-coding regulatory RNA molecules and their cognate mRNA targets. Indeed, in recent work, we revealed that Hfq condensates contribute to the stability of non-coding RNA and repression of sugar uptake via the phosphotransferase system during N starvation (17).

**Fig 1.**
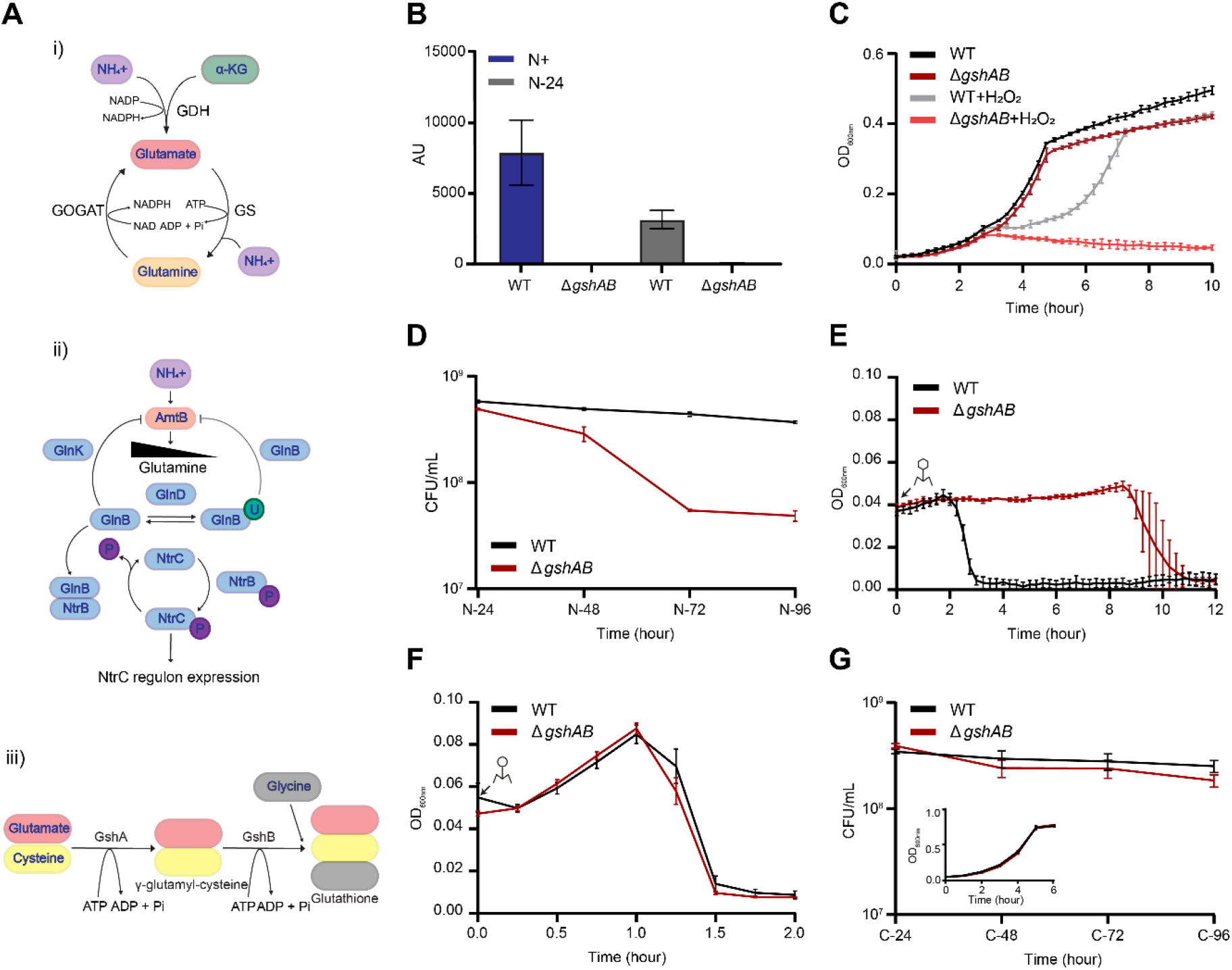
A specific role for GSH in the adaptive response to long-term N starvation. (A) i) Schematic showing the assimilation of ammonium and α-KG into glutamate and glutamine. ii) Schematic showing the cascade of signalling events that lead to the expression of the NtrC regulon under N starvation. iii) Schematic of the biosynthetic pathway that results in GSH synthesis. For i) to iii) see text for details. (B) Abundance of glutathione detected by targeted mass spectrometry in WT and Δ*gshAB E. coli* at N+ and N-24. (C) Growth measured by OD_600nm_ of WT and Δ*gshAB E. coli* during N starvation, with and without addition of H_2_O_2_ at N+. (D) Viability of WT and Δ*gshAB E*. coli during long-term N starvation measured by enumerating colony forming units. Cultures were sampled every 24-hours following N-24. (E) Collapse of WT and Δ*gshAB* N-24 *E. coli* cultures following infection with T7 phage. (F) as in (E), but for cultures of N+ *E. coli*. (G) Viability of WT and *ΔgshAB E. coli* during long-term C starvation measured by enumerating colony forming units. Inset shows growth of WT and Δ*gshAB E. coli* under C-limiting conditions. In (B)-(G), error bars represent standard deviation (n=3).

Glutamate, in addition to serving as a substrate for glutamine synthetase (for glutamine synthesis) also serves as a substrate for γ-glutamylcysteine synthetase (GshA), which catalyses the first step in the two-step pathway of glutathione synthesis (Fig. 1A(iii)). The second step is catalysed by glutathione synthetase (GshB). Glutathione (GSH) is an antioxidant tripeptide (γ-glutamyl-cysteinyl-glycine) that plays essential roles in *E. coli* cellular defense and metabolism, including protecting cells against oxidative stress by scavenging reactive oxygen species, maintaining the cellular redox balance, and acting as a cofactor for various enzymes; GSH also participates in the detoxification of harmful compounds (heavy metals) and helps maintain protein thiols in their reduced state (reviewed in (18)). Although GSH has a role in the adaptive response to various stresses and its intricate link to N metabolism (Fig. 1A(i-iii), the role of GSH in the adaptive response to N starvation in *E. coli* is unknown. By characterising an *E. coli* mutant unable to synthesise GSH (Δ*gshAB*) under N starvation, we show that GSH has pleiotropic, yet mutually exclusive, functions in N starved *E. coli*, adversely affecting survival, growth recovery and the temporal dynamics of Hfq condensation.

## RESULTS

### GSH has a specific role in the adaptive response to long-term N starvation

To characterise the Δ*gshAB* mutant bacteria under N starvation, we grew a batch culture of *E. coli* strain MG1655 in a highly defined minimal growth medium with a limiting amount of ammonium chloride as the sole N source (19). Under these conditions, when ammonium chloride in the growth medium runs out (N-), the bacteria enter a state of N starvation and become growth attenuated. In control experiments, we used liquid chromatography-electrospray ionization mass spectrometry to confirm that GSH was indeed not present in Δ*gshAB* bacteria during active growth under N replete conditions (N+) and in Δ*gshAB* bacteria that have been starved for 24 hours (N-24) (Fig. 1B). Hydrogen peroxide (H_2_O_2_) is a major contributor to oxidative damage and is effectively neutralised by GSH. As shown in Fig. 1C, the addition of H_2_O_2_ at N+ compromised, but did not fully inhibit, the growth of WT bacteria. However, H_2_O_2_ addition adversely affected the growth of Δ*gshAB* bacteria, affirming the importance of GSH in defense against oxidising agents (Fig. 1C) and the absence of GSH in Δ*gshAB* bacteria (Fig. 1B). In the absence of H_2_O_2_, the growth dynamics of Δ*gshAB* bacteria did not markedly differ from that of wildtype (WT) bacteria during N replete conditions and both strains became growth attenuated around the same time, suggesting that dynamics of N assimilation is unaffected in Δ*gshAB* bacteria (Fig. 1C). At N-24, the proportion of viable cells in the N-24 WT and Δ*gshAB* population did not differ (Fig. 1D). However, as N starvation persisted, the proportion of viable cells in the Δ*gshAB* population began to decline and by N-96h only ∼13% of cells in the Δ*gshAB* population were viable compared to those in the WT population (Fig. 1D). The compromised viability of *E. coli* during prolonged N starvation suggests that the adaptive metabolism is perturbed when the bacteria cannot synthesise GSH. The efficacy by which bacteriophages infect and replicate in bacteria can serve as an indicator of bacterial metabolic ‘health’. Therefore, we measured how quickly the prototypical *E. coli* bacteriophage T7 replicated and caused collapse of the N-24 WT and Δ*gshAB* cultures. As shown in Fig. 1E, following addition of T7, the culture of N-24 Δ*gshAB* bacteria collapsed ∼6.5h after that of the WT bacteria. This lag in culture collapse was not seen when N+ WT and Δ*gshAB* cultures were infected with T7 (Fig. 1F). To understand whether the long-term starvation survival defect of Δ*gshAB* bacteria is specific to N starvation, we conducted experiments in growth media with excess N source but containing limiting glucose (the sole carbon (C) source). Therefore, growth attenuation in this media correlates with C starvation. As shown in Fig. 1G (*inset*), the growth dynamics of Δ*gshAB* bacteria did not markedly differ from that of WT bacteria during C replete conditions. Notably, unlike under N starvation, the survival dynamics of Δ*gshAB* and WT bacteria were similar during prolonged C starvation (Fig. 1G). We conclude that GSH has a specific role in the adaptive response to long-term N starvation in *E. coli*.

### GSH regulates the temporal dynamics of Hfq condensation during N starvation

We previously reported that Hfq condensation is a hallmark subcellular response to long-term N starvation as they are not detected at N-but form progressively as N starvation ensues, becoming detectable ∼6 hours into N starvation. Given that Δ*gshAB* bacteria display compromised survival under long-term N starvation, we used Hfq condensation as a proxy for perturbed cellular adaptation (see Introduction) and monitored the dynamics of Hfq condensation by photoactivated localization microscopy (PALM) combined with single-molecule tracking of individual Hfq molecules in WT and Δ*gshAB* bacteria during N starvation. As a quantitative parameter to measure Hfq condensation we calculated the proportion of total Hfq molecules with an apparent diffusion less than 0.08 μm^2^/s, which had previously been defined as the ‘immobile’ population of Hfq molecules (%H_IM_) (16). The %H_IM_ was calculated based on all trajectories of Hfq in 50-200 bacterial cells within a given field of view. As shown in Fig. 2A, we did not detect any differences in %H_IM_ between WT and Δ*gshAB* bacteria at N+. However, upon N run-out (N-) and as N starvation set in for a prolonged period (N-, N-24), the %H_IM_ in Δ*gshAB* bacteria was consistently higher, with condensates clearly detectable within ∼3 hours under N starvation in Δ*gshAB* bacteria, indicative of increased Hfq condensation, than in WT bacteria. To establish that the observed differences in Hfq condensation is linked to GSH, we exogenously added 1 mM GSH to both WT and Δ*gshAB* bacteria at N+ and monitored Hfq condensation by PALM. As shown in Fig. 2B, exogenous addition of GSH reverted the temporal dynamics of Hfq condensation in Δ*gshAB* bacteria to that seen in WT bacteria. We note that the temporal dynamics of Hfq condensation in WT bacteria were highly similar between GSH untreated (Fig. 2A) and treated (Fig. 2B) cells. In previous work, we showed that Hfq condensates are absent in *E. coli* that have been specifically C starved for 24 hours (C-24). Therefore, to investigate whether the enhanced propensity to form condensates is general property Δ*gshAB* bacteria or, like in WT bacteria, a specific response to long-term N starvation, we compared the Hfq condensation dynamics in WT and Δ*gshAB* C-24 bacteria. As shown in Fig. 2C, we failed to detect distinct Hfq condensates in C-24 WT and Δ*gshAB* C-24 bacteria. We conclude that GSH is a regulator of Hfq condensation, and its absence perturbs the temporal dynamics of Hfq condensation, which is indicative of perturbed cellular adaptation during prolonged N starvation.

**Fig 2.**
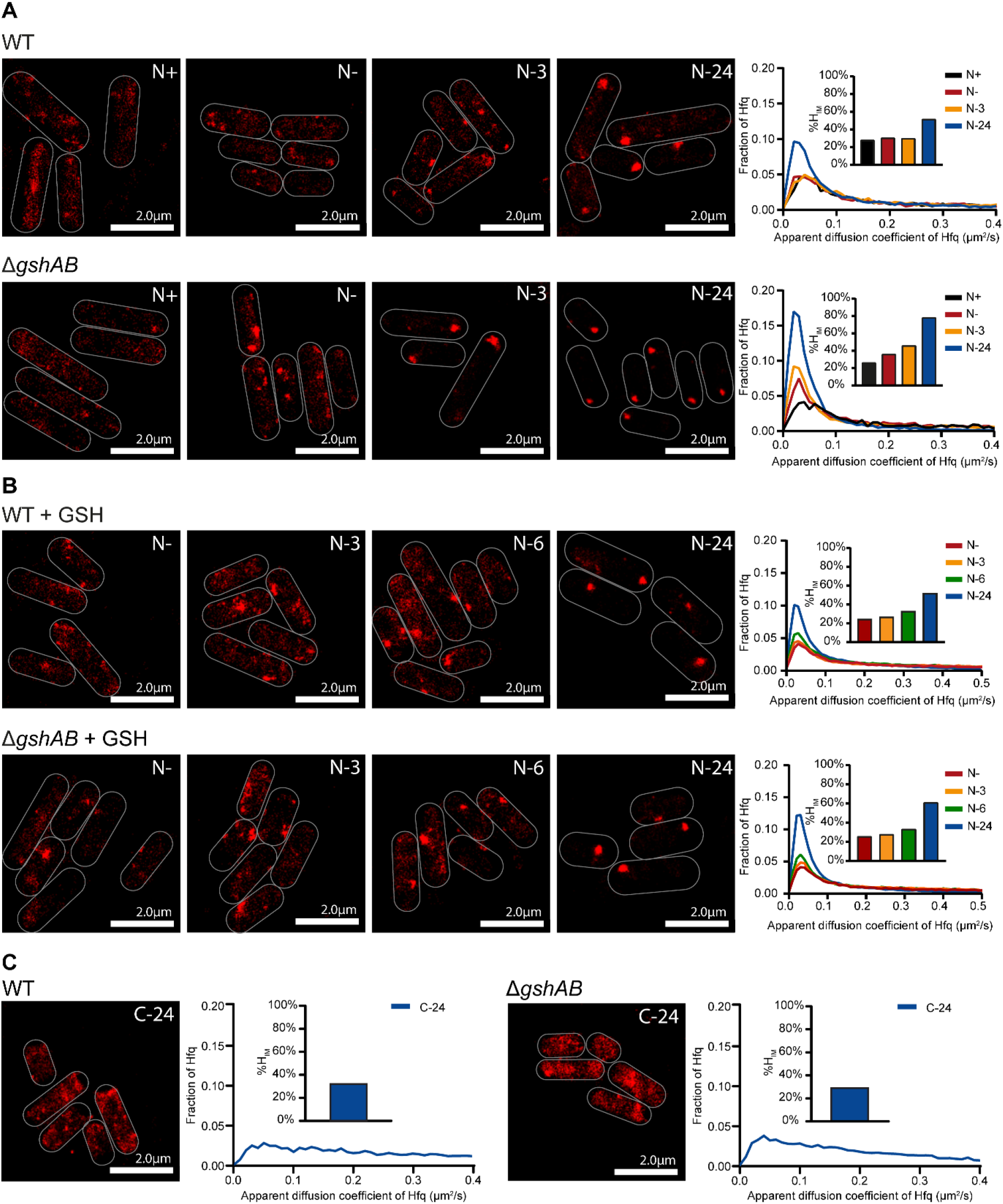
The regulation of the temporal dynamics of Hfq condensation by GSH during N starvation (A) Representative PALM images of Hfq in WT (top) and Δ*gshAB* (bottom) *E. coli* as a function of time during N starvation. Images taken at indicated timepoints. Graphs show the apparent diffusion coefficient of Hfq, inset bar graphs show corresponding %H_IM_ values. (B) as in (A) but when 1mM of GSH is added at N+. (C) as in (A) but for 24h C starved WT and Δ*gshAB* bacteria (C-24)

### The properties of Hfq condensates that form in WT and Δ*gshAB* bacteria are alike

We previously reported that Hfq condensation during long-term N starvation is dependent on the PTS regulator TmaR and that Hfq condensates form by a process analogous to liquid-liquid phase separation (LLPS)(15). To study the properties of the Hfq condensates that form in Δ*gshAB* bacteria, we monitored Hfq condensation dynamics in Δ*gshAB* bacteria devoid of TmaR (Δ*gshAB*Δ*tmaR*) and the sensitivity of the Hfq condensates in Δ*gshAB* bacteria to hexanediol (HEX), an aliphatic alcohol that disrupts LLPS condensates. Initially, however, we wanted to exclude the possibility that the altered Hfq condensation dynamics in Δ*gshAB* bacteria was not due to any aberrant increase Hfq protein levels. Indeed, as shown in Fig. 3A, the Hfq protein levels in WT and Δ*gshAB* bacteria were very similar at N+, N- and, importantly, at N-3, when the difference in Hfq condensation is detected between WT and Δ*gshAB* bacteria. As in Δ*tmaR* bacteria, we did not detect any Hfq condensates in N-24 (Δ*gshAB*Δ*tmaR*) bacteria (Fig. 3B), suggesting that TmaR remains a requirement for Hfq condensation in Δ*gshAB* bacteria. Further, the Hfq condensates in N-24 WT and Δ*gshAB* bacteria were both sensitive to HEX and reformed upon removal of HEX, suggesting that the altered temporal Hfq condensation dynamics in Δ*gshAB* bacteria was not due to any aberrant aggregation of Hfq (Fig. 3C). We conclude that the properties of Hfq condensates in WT and Δ*gshAB* bacteria are alike.

**Fig 3.**
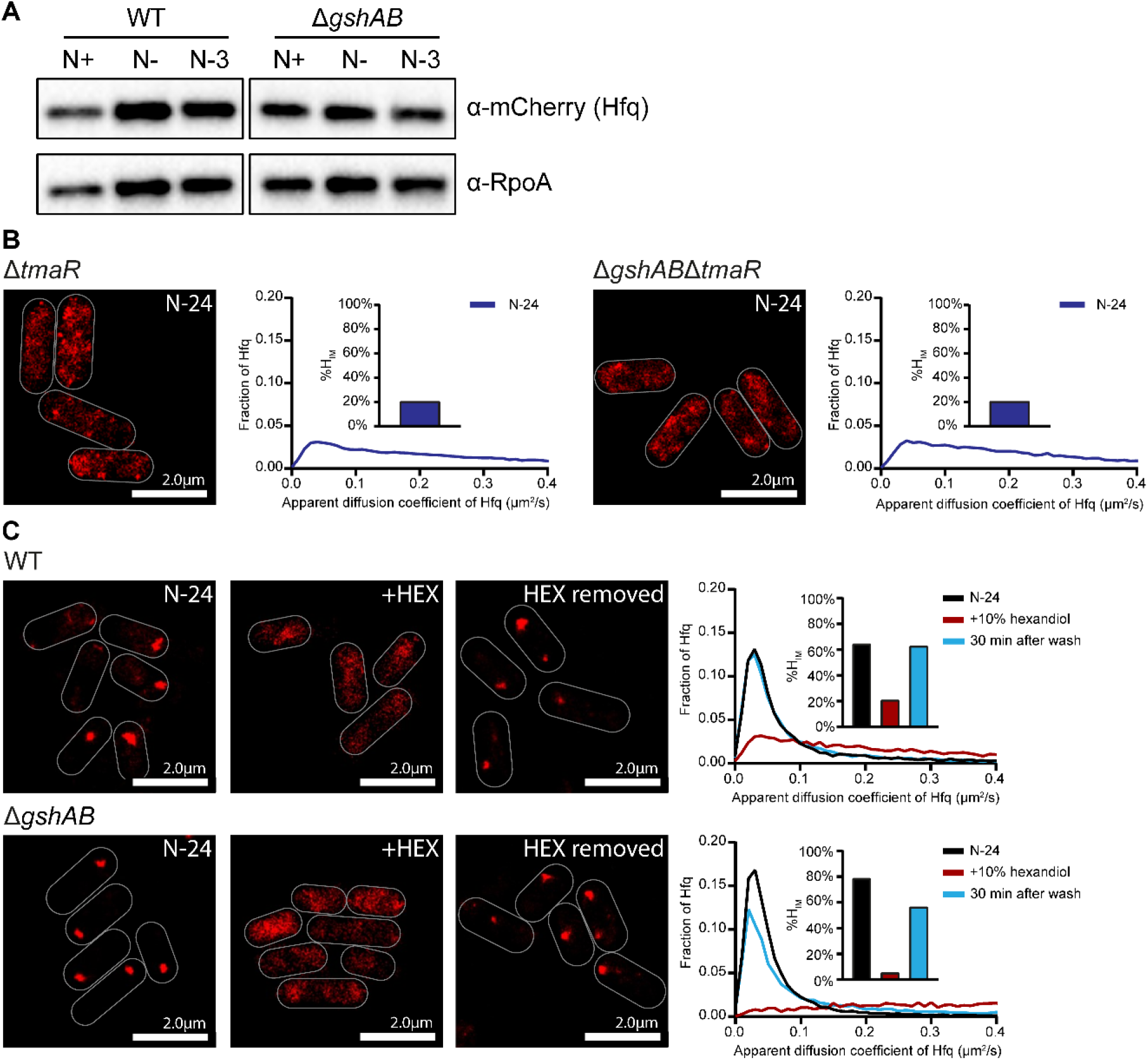
Hfq condensates in WT and Δ*gshAB* bacteria are alike. (A) Representative immunoblot of whole cell extracts of WT and Δ*gshAB E. coli* with Hfq translationally fused with PAmCherry, sampled at N+, N- and N-3 and probed with anti-mCherry antibody (for Hfq) and anti-RpoA antibody (loading control). (B) Representative PALM images of Hfq in Δ*tmaR* (left) and Δ*tmaR*Δ*gshAB* (right) *E. coli* at N-24. Graphs show the apparent diffusion coefficient of Hfq, inset bar graphs show corresponding %H_IM_ values. (C) as in (B), but for WT and Δ*gshAB E. coli* at N-24, following treatment with 10% hexanediol (HEX), and 30min following removal of HEX.

### GSH does not regulate the temporal dynamics of Hfq condensation by serving as an alternative N source in N starved *E. coli*

We observed that when GSH was added at N+, bacteria in the treated culture only became growth attenuated ∼10h after inoculation, whereas the bacteria in the untreated culture, as expected, entered growth attenuation at N-(∼5h after inoculation under our conditions) (Fig. 4A). We thus considered whether GSH is used as an alternative N source by N starved *E. coli* when the main N source, ammonium chloride, has run out. We deleted *ggt*, the gene which encodes γ-glutamyl transpeptidase, responsible for breaking down extracellular GSH, in the Δ*gshAB* strain and compared the growth and Hfq condensation dynamics in the WT, Δ*gshAB* and Δ*gshAB*Δ*ggt* bacteria during N starvation. As shown in Fig. 4B, the exogenous addition of GSH at N+ still simulated growth of Δ*gshAB*Δ*ggt* bacteria, as observed with WT and Δ*gshAB* bacteria (Fig. 4B), suggesting that it is unlikely that GSH is used as an alternative N source. However, the temporal dynamics of Hfq condensation did not differ between the Δ*gshAB* and Δ*gshAB*Δ*ggt*, with Hfq still forming condensates earlier during N starvation in Δ*gshAB* and Δ*gshAB*Δ*ggt* compared to in WT bacteria (compared Fig. 4C and Fig. 2A). Strikingly, the exogenous addition of GSH to Δ*gshAB*Δ*ggt* bacteria reverted the temporal dynamics of Hfq condensation to that seen in WT bacteria (Fig. 4D), demonstrating that the addition of exogenous GSH does not require its breakdown to revert condensate formation dynamics back to that of WT bacteria. We conclude that GSH does not influence the temporal dynamics of Hfq condensation by serving as an alternative N source during N starvation. Additional control experiments confirmed this conclusion: When aspartate, a poorer N source than ammonium chloride, was added to Δ*gshAB* bacteria at N+, we did not detect the reversion of the temporal dynamics of Hfq condensation to that seen with GSH treated cells (compare Fig. 4D with Fig. 4E), despite the aspartate treated cells becoming growth attenuated ∼5h after untreated bacteria (Fig. 4F). Of note, as aspartate contains only one N whereas GSH contains three, we added three times the concentration of aspartate than glutathione.

**Fig 4.**
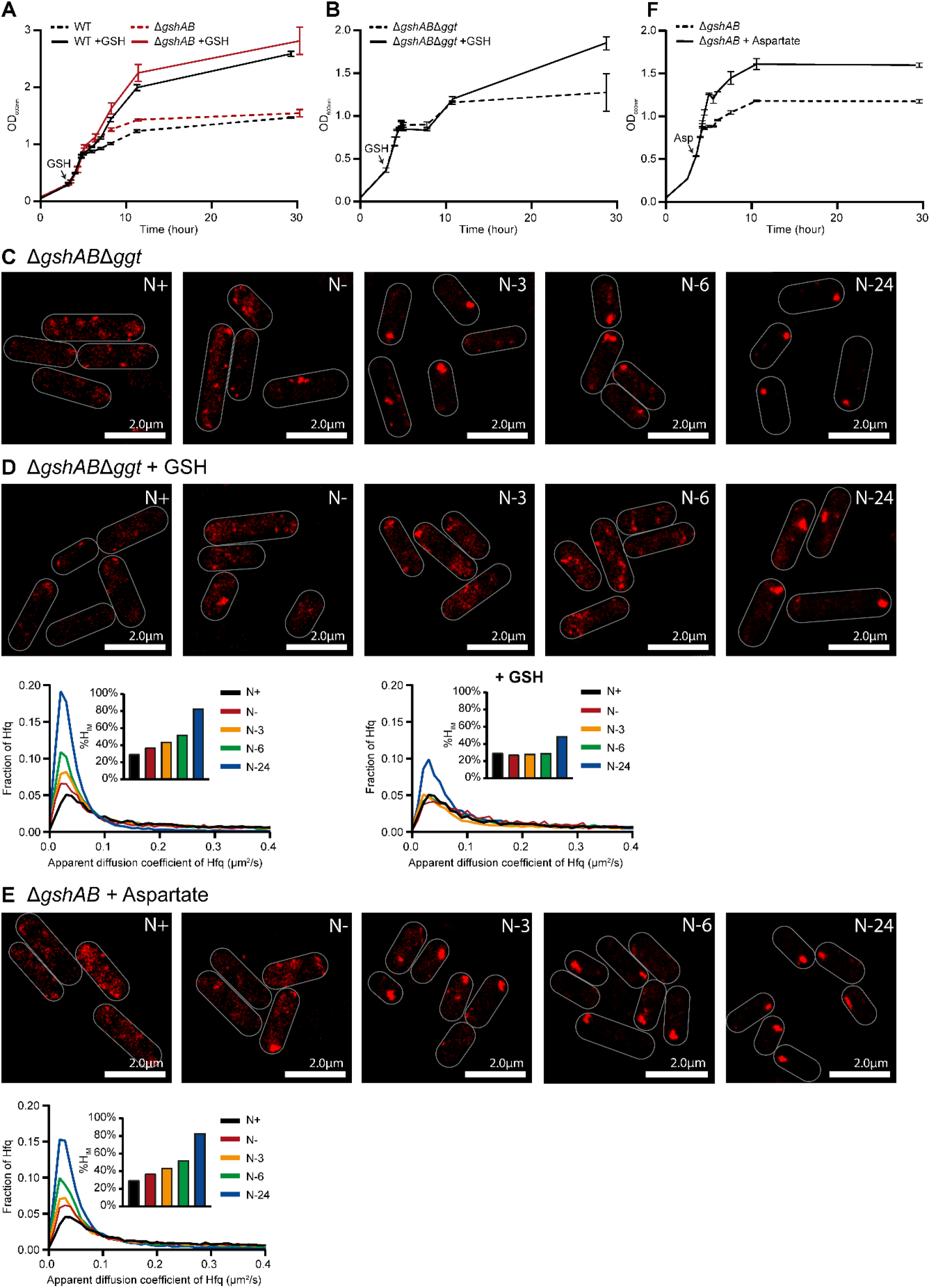
GSH does not affect Hfq condensation dynamics by serving as an N source. (A) Growth of WT and Δ*gshAB E. coli*, with and without addition of 1mM GSH at N+, as measured by OD_600nm_. Error bars represent standard deviation (n=3). (B) as in (A), but of Δ*gshAB* and Δ*gshAB*Δ*ggt E. coli*. (C) Representative PALM images of Hfq in Δ*gshAB*Δ*ggt E. coli* as a function of time during N starvation. (D) as in (C), but with addition of 1mM GSH at N+. Graphs show the apparent diffusion coefficient of Hfq, inset bar graphs show corresponding %H_IM_ values for (C) (Left) and (D) (Right). (E) as in (C), but for Δ*gshAB E. coli* to which 3mM aspartate was added at N+. (F) as in (A), but for Δ*gshAB E. coli* to which 3mM aspartate was added at N+.

### The absence of GSH compromises the growth recovery from N starvation

The addition of an N source to N-24 bacteria causes the dispersion of the Hfq condensates and growth resumption. Notably, the Hfq condensates also disperse when N-24 bacteria are resuspended into fresh media containing only N but not C and growth thus cannot resume, suggesting that Hfq condensates specifically respond to the N status of the cell and do not correlate with growth recovery *per se* (16). To better understand the how GSH influences Hfq condensation, we monitored the temporal dispersion dynamics of Hfq condensates and growth recovery of Δ*gshAB* bacteria. As shown in Fig. 5A, and as expected, the resuspension of N-24 WT bacteria into growth-permissive media (replete of N and C), led to the dispersion of Hfq condensates and ∼3 hours after resuspension the Hfq condensates were fully dispersed. In contrast, this was not the case when Δ*gshAB* bacteria were resuspended into growth-permissive media. As shown in Fig. 5B, even after ∼10 hours following resuspension, the Hfq condensates were present in Δ*gshAB* bacteria, but were eventually dispersed by ∼13 hours after resuspension. Control experiments showed that the dispersal of the Hfq condensates in Δ*gshAB* bacteria was still dependent on the N status (Fig. 5C) like in WT bacteria (16). Notably, the growth recovery of WT and Δ*gshAB* bacteria in growth-permissive media significantly differed and appeared to correlate with the dispersion of the Hfq condensates. As shown in Fig. 5D, the lag time to growth recovery (t_lag_) was ∼3 hours for WT bacteria and ∼10 hours for Δ*gshAB* bacteria. To investigate whether the dispersion of the Hfq condensates indeed correlates with growth recovery, we conducted experiments with Δ*gshAB*Δ*tmaR* bacteria in which Hfq condensates do not form (Fig. 3B). As shown in Fig. 5E, the t_lag_ between WT and Δ*tmaR* bacteria differed by ∼2.6 hours but this difference increased to ∼17.1 hours in the Δ*gshAB* background (i.e. Δ*gshAB*Δ*tmaR* bacteria). Consistent with a specific role for GSH in adaptive response to long-term N starvation (see Fig. 1), the t_lag_ between C-24 WT and Δ*gshAB* bacteria only differed by ∼1.44 hours (Fig. 5F). We conclude that the inability of *E. coli* to synthesise GSH compromises its ability to recover specifically from long-term N starvation. We further conclude that the compromised growth recovery and the altered dispersion dynamics of Hfq condensation are mutually exclusive properties of N starved *E. coli* devoid of GSH.

**Fig 5.**
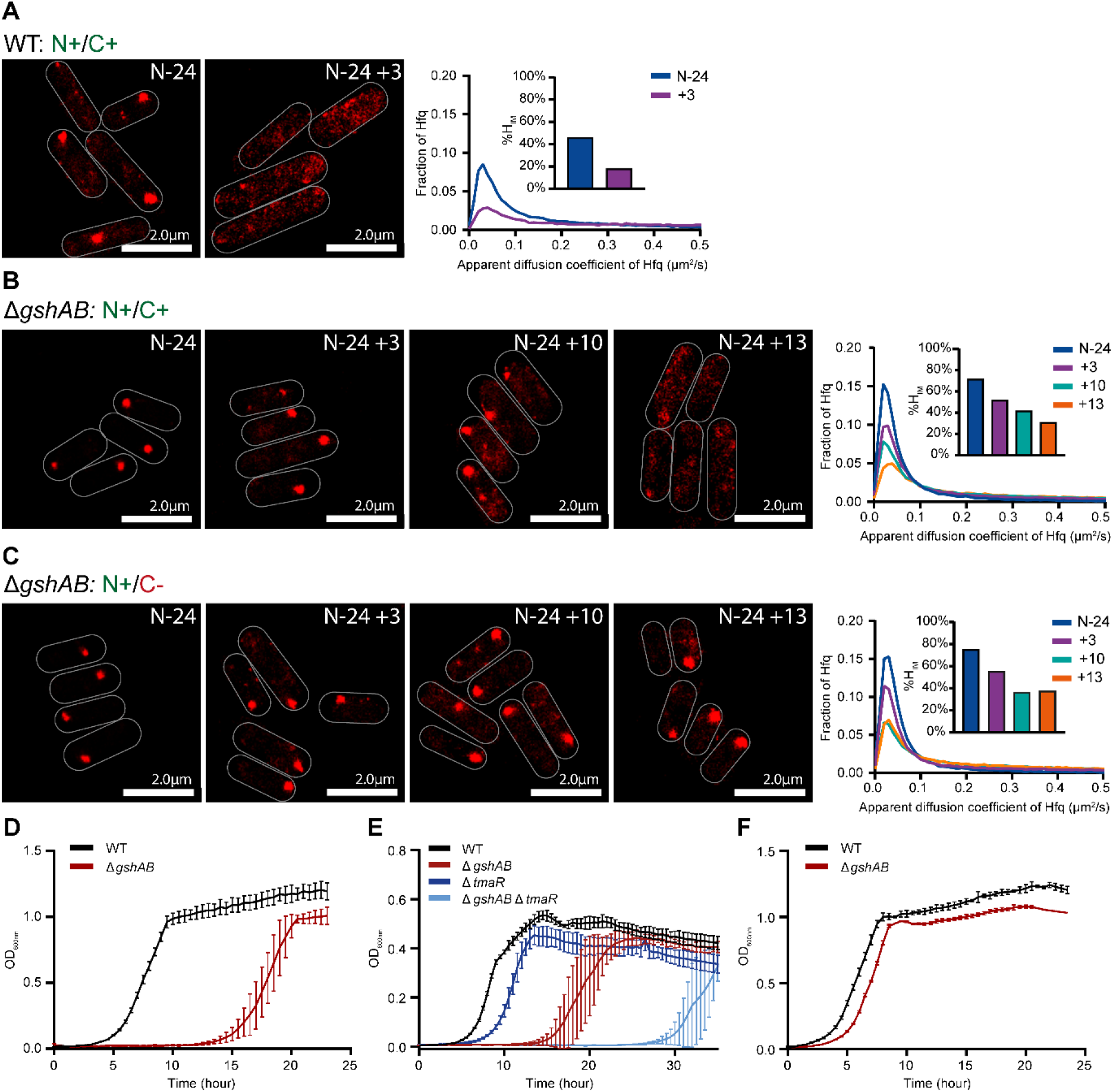
GSH-deficient bacteria experience compromised growth recovery. (A) Representative PALM images of Hfq in WT *E. coli* at N-24 and 3 hours following resuspension into growth-permissive media (N+/C+). Graph shows the apparent diffusion coefficient of Hfq, inset bar graph show corresponding %H_IM_ values. (B) as in (A) but for Δ*gshAB E. coli* imaged 3, 10 & 13 hours post resuspension into growth-permissive media. (C) as in (B) but for Δ*gshAB E. coli* resuspended into fresh media with N, but without C (N+/C-). (D) Growth-recovery of WT and Δ*gshAB E. coli* from N-24 following subculturing into growth-permissive media. Error bars represent standard deviation (n=3). (E) as in (D), but for WT, Δ*gshAB*, Δ*tmaR* and Δ*gshAB*Δ*tmaR E. coli*. (F) as in (D), but for WT and Δ*gshAB E. coli* initially grown to C-24 and subsequently subcultured into growth-permissive media.

## DISCUSSION

Bacterial adaptive responses to environmental stresses are often investigated under acute exposure conditions. Consequently, the metabolic and gene expression changes that underpin adaptation during prolonged stress remain poorly understood. In *E. coli* and related bacteria, GSH is amongst the most abundant metabolites, primarily functioning as an antioxidant and detoxifying agent (18). *E. coli* strains lacking GSH exhibit normal growth rates even in minimal media yet display heightened sensitivity to oxidative stress (see Fig. 1C). Through characterization of an *E. coli* mutant deficient in both GSH biosynthetic enzymes (Δ*gshAB*), we have uncovered pleiotropic roles for GSH in the adaptive mechanisms that support survival under prolonged N starvation. Notably, GSH deficiency does not impair adaptation to prolonged C starvation, highlighting a specific requirement for GSH in the N starvation adaptive response.

Biomolecular condensates have emerged as a key mechanism for subcellular organization in bacteria, enabling spatial and temporal regulation of cellular processes (20-22). Among these, condensates formed by Hfq are a hallmark feature of stress-associated biomolecular assemblies (15-17). During N starvation, Hfq condensates progressively appear as the stress intensifies. Our new findings reveal that in *E. coli* cells lacking GSH, Hfq condensates form significantly earlier, suggesting that GSH deficient cells enter a heightened stress state more rapidly than their WT counterparts. Given GSH’s well-established antioxidant properties, it is widely used as an anti-ageing agent in the cosmetic industry. Thus, this accelerated Hfq condensation may reflect a form of premature cellular ageing specifically in GSH deficient, N-starved bacteria. For example, proteins such as MetE (methionine biosynthesis) and DnaK (protein quality control), are known to be post-translationally modified via glutathionlylation (i.e. the binding of GSH to cysteine residues), which protects them from permanent damage (23). Thus, premature ageing of cells lacking GSH could be due to irreversible damage to key proteins. Further, building on our previous findings that Hfq condensates can be induced by the exogenous addition of α-KG (17), our new results suggest that the condensation dynamics of Hfq (and potentially other proteins) are modulated by metabolite fluxes during stress adaptation. These observations underscore the possibility that specific metabolites, act as cues and influence the formation and timing of biomolecular condensates in response to stresses to spatiotemporally organise and concentrate specific cellular processes in the stressed cell.

The heterotypic composition of biomolecular condensates and the pleiotropic nature of the proteins involved in their formation make it challenging to assign discrete functions to these assemblies. Hfq condensates, which emerge progressively during N starvation in *E. coli* and disperse upon N replenishment. Therefore, it is conceivable that they are likely to play roles in both adaptation to nutrient stress and recovery. Indeed, our findings appear to show that in GSH-deficient bacteria, Hfq condensates disperse significantly more slowly than in WT cells, and that the timing of dispersal coincides with the recovery of growth following N starvation. However, this correlation does not imply causality. In fact, our data suggest that Hfq condensates do not directly contribute to growth recovery. Rather, consistent with our previous results (17), they appear to contribute to post-transcriptional and metabolic regulation during adaptation to prolonged N starvation *per se*.

Our findings suggest a specific role for GSH in facilitating the growth recovery of N-starved bacteria. Given GSH’s multifaceted involvement in bacterial cellular processes and stress responses (18), pinpointing its exact contribution to recovery is challenging. Notably, GSH can be catabolized by γ-glutamyltransferase (GGT) into cysteine and glycine, thereby replenishing the intracellular amino acid pool (24). It is therefore plausible that, under N starvation, changes in amino acid availability may be responsible for the demise of and impaired recovery in GSH-deficient bacteria during prolonged N starvation. However, since bacteria lacking GGT (Δ*ggt*) recovered like WT bacteria (Supplementary Fig. 1), this mechanism is unlikely to account for the compromised recovery phenotype of GSH deficient bacteria. GSH is also important for aconitase function, the enzyme that converts citrate to isocitrate, the first step of the Krebs cycle. Thus, the absence of GSH induces oxidative damage to aconitase (25-27), which could cause the buildup of citrate, which could differentially impair metabolite flux through the Krebs cycle during growth recovery (from N and C starvation). Indeed, citrate levels in N-24 Δ*gshAB* bacteria are ∼2.5-fold higher than in WT bacteria (Supplementary Fig. 2).

In sum, our new results have uncovered multiple, yet mutually exclusive, roles for GSH in the adaptive response to N starvation that potentially extends its canonical function as a stress protectant. Future research, focused on the temporal glutathionylation dynamics of proteins at a systems-wide scale and targeted measurement of metabolites of the Krebs cycle in WT and Δ*gshAB* bacteria during N and C starvation will uncover the precise mechanistic basis underpinning GSH’s functions in the adaptive response to N starvation.

## MATERIALS AND METHODS

### Bacterial strains and plasmids

All strains used in this study were derived from *Escherichia coli* K-12 and are listed in Supplementary Table 1. Gene deletions were introduced into the WT and Hfq-PAmCherry strains as described previously (11). Briefly, the knockout alleles were transduced using the P1*vir* bacteriophage with strains from the Keio collection (28) serving as donors. Where multiple modifications were introduced into a strain, the existing *kanR* cassette was first cured by expressing the yeast *flp* flippase recombinase from pCP20 (29).

### Bacterial growth conditions

N starvation experiments were carried out as previously described in (19). Briefly, unless otherwise stated, bacterial cultures were grown in Gutnick minimal medium (33.8 mM KH_2_PO_4_, 77.5 mM K_2_HPO_4_, 5.74 mM K_2_SO_4_, 0.41 mM MgSO_4_) supplemented with Ho-LE trace elements (30), 0.4% (w/v) and 10 mM NH_4_Cl (for overnight cultures and recovery experiments) or 3 mM NH_4_Cl (for day cultures) at 37°C in a shaking (180 rpm) incubator. Bacterial day cultures for C starvation experiments were grown in Gutnick minimal media supplemented with 10mM NH_4_Cl and 0.06% (w/v) glucose. For experiments containing HEX, HEX was added at 10% w/v and cells imaged on agarose slides containing 5% HEX. For GSH addition experiments, GSH was added to a final concentration of 1mM at N+. The proportion of viable cells in the bacterial population was determined by enumerating CFU/ml from serial dilutions on lysogeny broth agar plates.

### T7 phage infection assay

Bacterial cultures were grown in Gutnick minimal medium as described above to the indicated time points. Bacterial culture samples were taken, centrifuged and resuspended in fresh Gutnick minimal media supplemented with either 2mM NH_4_Cl and 12.5mM glucose (for N+) or 5mM glucose (for N-24) and diluted to *A*_600 nm_ of 0.3 to a final volume of 500 μl and transferred to a flat-bottomed 48-well plate, together with T7 phage at a final concentration of 4 × 10^9^ phage/ml. The cultures were then grown at 37°C with shaking at 700 rpm in a SPECTROstar Nano microplate reader (BMG LABTECH), and *A*_600 nm_ readings were taken every 10 min.

### Targeted metabolite measurement

At the indicated time point, approximately 10^10^ cells were collected, and washed twice in ¼ strength Ringer’s solution (Thermo Scientific, BR0052G). Cell pellets were resuspended in 500μl of cold (-20°C) methanol:acetonitrile:water (2:2:1, v/v/v) + 0.1% formic acid. Samples were stored at -80°C until analysis. Before analysis, all vials received were reconstituted with 150 µL of 97.5% H_2_O + 2.5% acetonitrile + 0.2% formic acid (FA), diluted, vortexed and transferred to inserts. Pooled quality control of all samples was then generated by pooling 10 µL of the first replicate for each experimental condition and injected every 8 samples. All reagents used were of ultra-high-performance liquid chromatography (UHPLC) gradient grade and all standards were of analytical grade. Targeted metabolomics analysis was performed using an Agilent 1290 Liquid chromatography (LC) system (Agilent Technologies, CA, USA) coupled to a QTRAP 4000 mass spectrometry (MS) system (SCIEX, Danaher, WA, USA). Chromatographic separation was achieved using a Luna Omega Polar C18 column (Phenomenex/Danaher, WA, USA). The analysis was conducted in positive ion mode (A: H2O + 0.2% FA / B: acetonitrile + 0.2% FA) on a 20-minute gradient and in negative ion mode (A: H2O + 0.1% FA + 10 mM ammonium formate / B: 100% acetonitrile) on a 14-minute gradient at 0.450 ml/min flow rate. All data was acquired in multiple reaction monitoring (MRM) mode. Resulting spectra analysed using an in-house data analysis workflow based on (31).

### Immunoblotting

Immunoblotting was conducted in accordance with standard laboratory protocols, with primary antibodies incubated overnight at 4°C, and secondary antibodies incubated for 1 hour at room temperature The following antibodies were used: rabbit polyclonal anti-mCherry (Abcam ab167453) at 1:100 dilution, mouse monoclonal anti-RpoA (Biolegend, WP003) at 1:100 dilution, HRP goat anti-rabbit IgG (GE Healthcare NA934-1ML) at 1:10000 dilution and HRP Goat anti-mouse IgG (Biolegend, 405306) at 1:10000 dilution. ECL Prime Western blotting detection reagent (GE Healthcare, RPN2232) was used to develop the blots, which were analysed on the ChemiDoc MP imaging system.

### Photoactivated localization microscopy (PALM) and single molecule tracking (SMT)

For the PALM and single-molecular tracking (SMT) experiments, the Hfq-PAmCherry, and mutant derivative, reporter strains were used. Bacterial cultures were grown as described above and samples were taken at the indicated time points, then imaged and analysed as previously described (32,33). Briefly, 1 ml of culture was centrifuged, washed and resuspended in a small amount of Gutnick minimal medium supplemented with N and C concentrations that reflected the concentration contained in the media at the time point sampled. One μl of the resuspended culture was then placed on a Gutnick minimal medium agarose pad (1% (w/v) agarose, 1x Gutnick minimal medium supplemented with N and C concentrations that reflected the concentration contained in the media at the time point sampled). Cells were imaged on a PALM-optimized Nanoimager (Oxford Nanoimaging, https://oni.bio/nanoimager/) with 15 millisecond exposures, at 66 frames per second over 10,000 frames. Photoactivatable molecules were activated using 405 nm and 561 nm lasers. Fields-of-view typically consisted of 100-200 bacterial cells.

For SMT, the Nanoimager software was used to localize the molecules by fitting detectable spots of high photon intensity to a Gaussian function. The Nanoimager software SMT function was then used to track individual molecules and draw trajectories of individual molecules over multiple frames, using a maximum step distance between frames of 0.6 μm and a nearest-neighbour exclusion radius of 0.9 μm. The software then calculated the apparent diffusion coefficients (*D\**) for each trajectory over at least four steps, based on the mean squared displacement of the molecule. To calculate %H_IM_, we collated D* values from multiple fields of view and determined the proportion of D* values that fell into our previously defined immobile population (D* ≤0.08 μm/s^2^) (16).

## ACKNOWLEDGMENTS

This work was supported by the Leverhulme Trust (RPG-2020-050) project grants to S.W and an MRC ICASE PhD studentship to H.E. Funding to pay the Open Access publication charges for this article was provided by Imperial Open Access Fund.

H.E, J.M and V.B. performed experiments and analysed the data.

H.E., J.M. and S.W. conceived and designed experiments and wrote the manuscript

