## Supplementary Figures for "Pleiotropic functions of glutathione in the adaptive response to long-term nitrogen starvation in *Escherichia coli*"

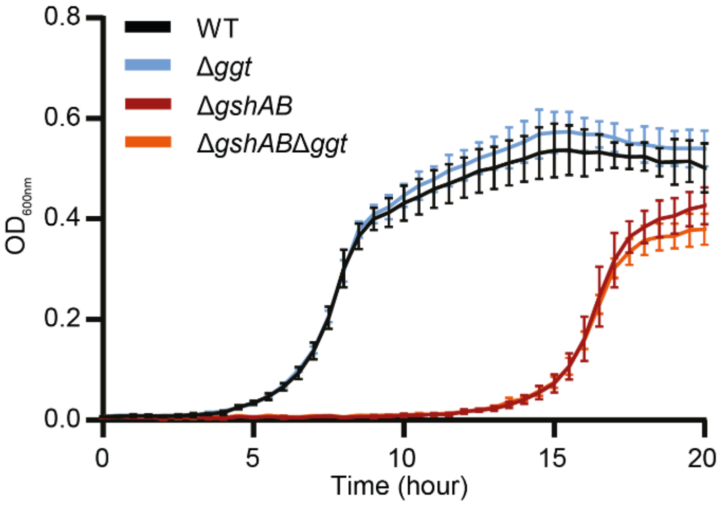


**Supplementary Fig 1.** Growth-recovery of WT, Δ*ggt*, Δ*gshAB* and Δ*gshAB*Δ*ggt* *E. coli* from N-24, following subculturing into growth-permissive media. Error bars represent standard deviation (n=3).

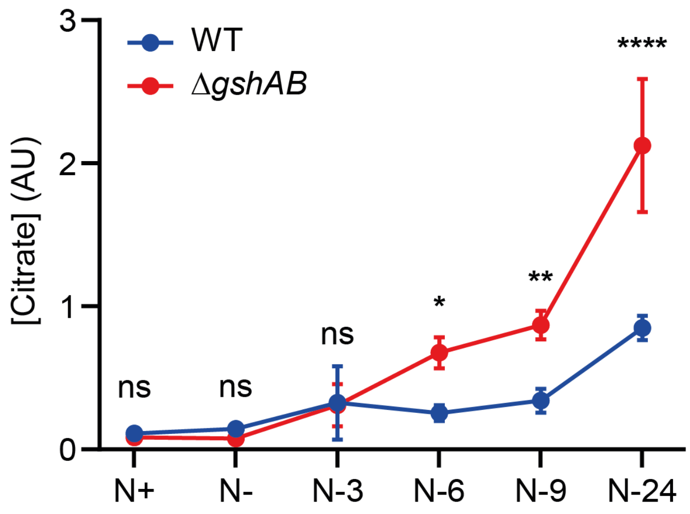


**Supplementary Fig 2.** Graph showing the intracellular concentration of citrate in WT (Blue) and Δ*gshAB* (Red) *E. coli* experiencing N starvation. Errors bars represent standard deviation (n=3). Statistical analysis performed by two-way ANOVA with Šidák multiple comparisons. (****, P<0.0001; **, P<0.01; *, P<0.05).

**Supplementary Table 1**

Strains used in this study

| Strain | Description | Source |
|---|---|---|
| WT MG1655 | *E. coli* K-12 *rph-1* | *E. coli* Genetic Stock Centre |
| Hfq-PAmCherry | MG1655 *hfq-PAmCherry-kan* | (1) |
| Δ*gshAB* | MG1655 Δ*gshAB::kan* | This work |
| Δ*gshAB* Hfq-PAmCherry | Hfq-PAmCherry Δ*gshAB::kan* | This work |
| Δ*tmaR* | MG1655 Δ*tmaR::kan* | This work |
| Δ*gshAB*Δ*tmaR* | MG1655 Δ*gshAB ΔtmaR::kan* | This work |
| Δ*gshAB*Δ*tmaR* Hfq-PAmCherry | Hfq-PAmCherry Δ*gshAB ΔtmaR::kan* | This work |
| Δ*ggt* | MG1655 Δ*ggt::kan* | This work |
| Δ*ggt* Hfq-PAmCherry | Hfq-PAmCherry Δ*gg*t::kan | This work |
| Δ*gshAB*Δ*ggt* | MG1655 Δ*gshAB Δggt::kan* | This work |
| Δ*gshAB*Δ*ggt* Hfq-PAmCherry | Hfq-PAmCherry Δ*gshAB Δggt::kan* | This work |

1. McQuail, J., Switzer, A., Burchell, L., and Wigneshweraraj, S. (2020) The RNA-binding protein Hfq assembles into foci-like structures in nitrogen starved Escherichia coli. *The Journal of biological chemistry* **295**, 12355-12367
